# Characterization of *Ralstonia pseudosolanacearum* diversity and screening host resistance to manage bacterial wilt in South Asia

**DOI:** 10.1101/2023.10.13.559983

**Authors:** Nagendra Subedi, Tabitha Cowell, Matthew Cope-Arguello, Pierce Paul, Gilles Cellier, Hashem Bkayrat, Nicolas Bonagura, Angela Cadatal, Rachel Chen, Ariana Enriquez, Rama Parasar, Lisa Repetto, Aracely Hernandez Rivas, Mahnoor Shahbaz, Kaitlin White, Tiffany M. Lowe-Power, Sally A. Miller

## Abstract

In South Asia, bacterial wilt pathogens in the *Ralstonia solanacearum* species complex (RSSC) impose major constraints on eggplant, tomato, and pepper production. To improve the efficacy of bacterial wilt management, the goals of this study were to (1) conduct a survey of RSSC pathogens in Bangladesh and Nepal, (2) characterize the genetic diversity of these isolates, and (3) screen 37 tomato, eggplant, and pepper accessions for resistance to six representative isolates from South Asia. We isolated 99 isolates from Bangladesh and 20 isolates from Nepal and determined that all are phylotype I isolates of the *Ralstonia pseudosolanacearum* species. We sequenced and assembled draft genomes for 25 isolates. Phylogenomic analyses suggest that there is a wide diversity of endemic phylotype I isolates in South Asia, and possible introductions of two clonal phylotype I lineages into Bangladesh and Nepal. We contextualize our newly described isolates based on prior reports of RSSC diversity in South Asia and global reports of RSSC pathogens on eggplant and pepper. Greenhouse trials revealed multiple tomato, eggplant, and pepper accessions that exhibit promising levels of resistance to six phylotype I isolates from South Asia.

## Introduction

Bacterial pathogens in the *Ralstonia solanacearum* species complex (RSSC) cause a group of related wilt diseases by colonizing xylem and impairing water transport (Kelman 1953). RSSC comprises economically significant pathogens of agronomically important crops, such as tomato, eggplant, potato, banana, peanut, ginger and others (Hayward 1991; Savary et al. 2019).

Bacterial wilt is a major constraint for production of eggplant (*Solanum melongena* also known as brinjal or aubergine), tomato (*S. lycopersicum*), and pepper (*Capsicum* spp.) in South Asia (Sinha, SK 1986; Sood and Singh 1993; Adhikari et al. 1993; Pradhanang et al. 2000; Pradhanang and Momol 2001; Ahmed et al. 2013; Adhikari et al. 1997; Singh et al. 2010). Host resistance is the most practical and sustainable approach for management of this disease, however very few bacterial wilt-resistant cultivars are available (López and Biosca 2005) (Pandiyaraj et al. 2019). The main sources of bacterial wilt resistance in tomato breeding populations are its wild relatives such as *S. pimpinellifolium*, *S. hirsutum* and *S. peruvianum* (Carmeille et al. 2006). Host resistance against bacterial wilt is strain-specific due to the considerable genetic diversity of the pathogen populations (Danesh and Young 1994; Wang et al. 1998; Lebeau et al. 2011). In the Check List of Commercial Varieties of Vegetables published by the Government of India, eight tomato, three eggplant, and no pepper varieties are listed as resistant to bacterial wilt (Singh 2012). The major bacterial wilt-resistant cultivars used in South Asia are tomato lines Arka Ananya, Arka Abhijit, Arka Abha, CLN2020C, All Rounder, Swarakhsha, Rakshak, and Trishul, and eggplant lines Kata Begun, Marich Begun, Pusa purple cluster, JC-2, Pant Samrat, Arka Anand, and Uttar (Dutta and Rahman 2012; Rahman et al. 2011; Singh 2012; Timila and Joshi 2007).

Grafting desired commercial varieties onto resistant rootstocks is another approach to combat bacterial wilt (Rivard and Louws 2011). Bacterial wilt-resistant *S. sisymbriifolium*, also known as sticky nightshade, fire-and-ice plant, litchi tomato, etc., is a popular rootstock in South Asia that is also resistant to *Meloidogyne* spp. nematodes that cause root-knot (Miller et al. 2005). Plants grafted onto *S. sisymbriifolium* not only reduce bacterial wilt incidence but also increase marketable yield, even in the absence of disease pressure. However, failures of *S. sisymbriifolium* resistance to bacterial wilt at several locations in Bangladesh and Nepal is a concern for researchers and growers in this region.

RSSC are classified into four phylotypes (I-IV) that emerged and diversified on different continents (Villa et al. 2005). Phylotype I emerged in continental Asia, II in the Americas, III in Africa, and IV in Indonesia/Southeast Asia (Villa et al. 2005). However, movement of plants through international trade has allowed phylotypes I and II strains to become widely established in new locations. The phylotypes are consistent with the division of RSSC into three species: *R. solanacearum* (phylotype II), *R. pseudosolanacearum* (phylotypes I and III), and *R. syzygii* (phylotype IV) (Prior et al. 2016; Safni et al. 2014). Recently, an international consortium of *Ralstonia* researchers reaffirmed that phylotypes I and III are two groups within the single *R. pseudosolanacearum* species (Lowe-Power et al. 2023).

Phylotype I is the most widespread phylotype in India and Sri Lanka (Ramesh et al. 2014; Gurjar et al. 2015; Sagar et al. 2014; Kumar et al. 2013, 2014; Ghorai et al. 2022), but phylotypes II and IV are also present in South Asia. Published studies and public genome databases indicate that isolates in the pandemic brown rot IIB-1 lineage are present as potato pathogens in India, Nepal, Bangladesh, and Sri Lanka (Pradhanang et al. 2000; Sagar et al. 2014; Gurjar et al. 2015; Cellier and Prior 2010). Additionally, phylotype IV has become established in the hills of Meghalaya, the Indian state east of Bangladesh (Gurjar et al. 2015; Sagar et al. 2014). Although RSSC are prevalent pathogens in Bangladesh and Nepal (Ahmed et al. 2013; Pradhanang et al. 2000; Hossain et al. 2022), little is known about their genetic diversity.

The objectives of this study were to characterize RSSC isolates from Bangladesh and Nepal and to screen a worldwide collection of tomato, eggplant and pepper genotypes against representative RSSC isolates from India, Bangladesh, and Nepal. In 2012, we purified 119 RSSC isolates from solanaceous crops in Bangladesh and Nepal and a representative subset of 25 isolates were sequenced for their genomes. We screened 37 plant accessions for bacterial wilt resistance, including the 30 accessions proposed as Core-TEP by Lebeau et al. (2011).

## Methods

### Bacterial isolates

We conducted a survey during 2012 to collect RSSC isolates in major vegetable growing regions of Bangladesh and Nepal (Fig. 1A and Table S1). Bacterial isolates were purified from symptomatic eggplant, tomato, pepper, potato (*Solanum tuberosum*), and *S. sisymbriifolium* (used as rootstock of tomato and eggplant scions) on CPG medium (1 g / L casamino acids, 10 g / L peptone, and 5 g / L glucose) with 1% tetrazolium chloride (Kelman 1954). The identity of isolates was confirmed as RSSC based on colony morphology, RSSC-specific ImmunoStrips (Agdia Inc., Elkhart, IN), and a polymerase chain reaction (PCR) assay using the RSSC-specific primers 759/760 (Opina et al. 1997) as described previously (Lewis Ivey et al. 2007). Six Indian isolates were also included in the study. All *Ralstonia* isolates were imported to Ohio under APHIS permit no. P526P-11-02092.

**Fig 1.**
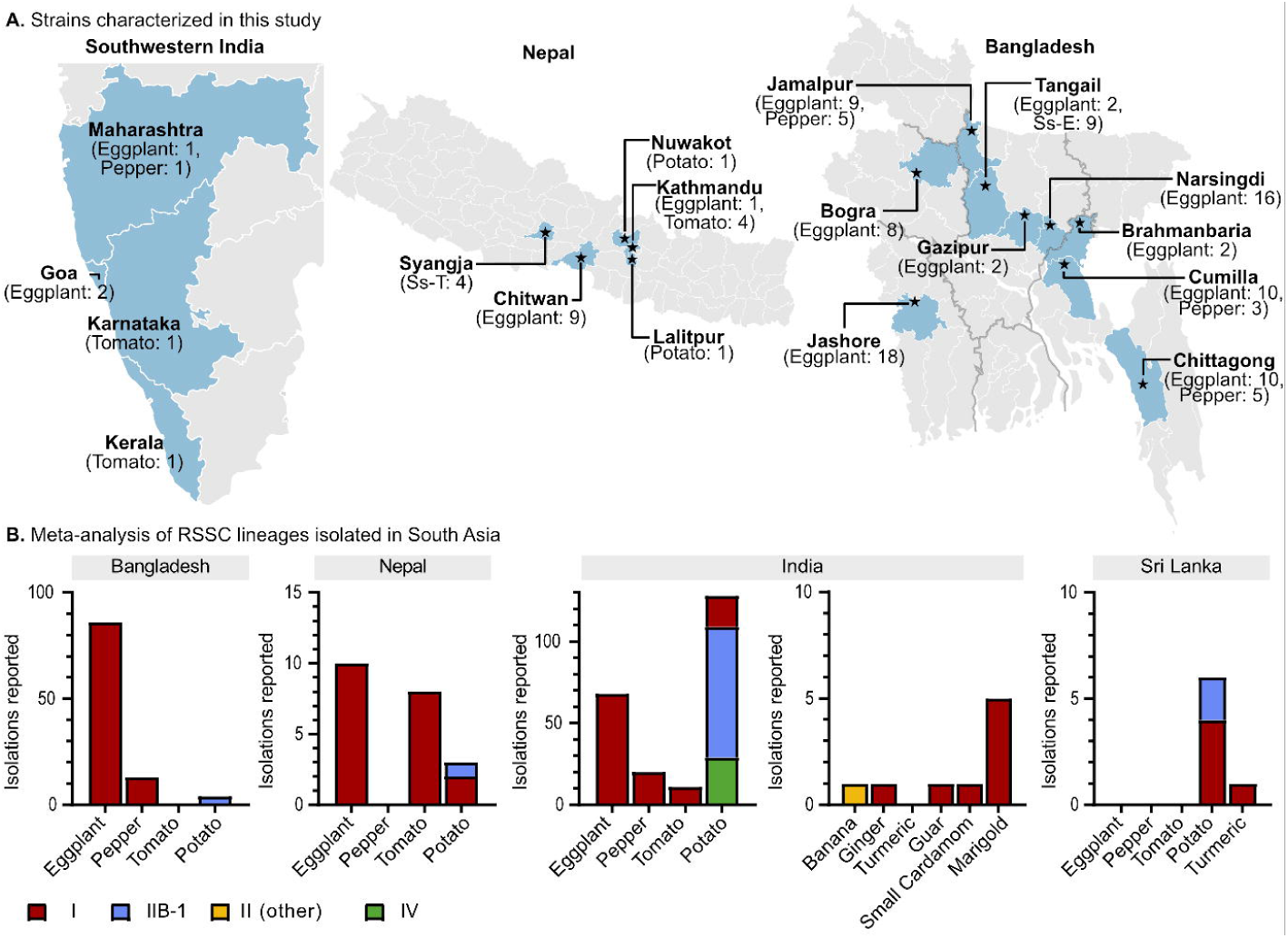
*Ralstonia solanacearum* species complex (RSSC) isolates in South Asia. (A) Origins of South Asian *R. pseudosolanacearum* phylotype I isolates characterized in this study. Stars indicate the sampling locations, and light blue shading indicates districts (Bangladesh/Nepal) and states (India) where isolates originated. Abbreviations: Ss-E, eggplant grafted on *S. sisymbriifolium* rootstock; Ss-T, tomato grafted on *S. sisymbriifolium* rootstock. (B) Meta-analysis of RSSC lineages isolated in South Asia in this study and the literature, adapted from Lowe-Power et al. 2022. This study included Bangladesh isolates (77 from eggplant, nine from grafted eggplant/*S. sisymbriifolium*, and 13 from pepper) and Nepal isolates (ten eggplant, four from tomato, two from potato, and two from grafted tomato/*S. sisymbriifolium*).

### Phylotype determination

Phylotype-specific multiplex PCR (Pmx-PCR) was performed using five phylotype-specific (Fegan and Prior 2005) and two species complex-specific primers (Opina et al. 1997). Reaction mixture preparation, amplification and gel electrophoresis were performed as described previously (Fegan and Prior, 2005; Lewis Ivey et al. 2007). Genomic DNA of isolates GMI1000, K60, UW386 and UW443 were used as positive controls for phylotypes I, II, III and IV respectively.

### Genome sequencing and assembly and quality control

Genomic DNA was extracted with Zymo Quick-DNA kits (Zymo Research, Irvine, CA). We used short-read Illumina sequencing to sequence draft genomes of twenty-four of the isolates. Library prep was performed using the Illumina DNA Prep kit (Illumina, Inc., San Diego, CA) following their standard gDNA library prep workflow. Nextera DNA CD Indexes (Illumina, Inc.) were used for indexing during library prep. The DNA input for each sample was within 100-500 ng, so quantification of the libraries was not performed and instead the library pooling protocol for DNA inputs of 100-500 ng was followed according to the manufacturer’s specifications. An aliquot of the pooled libraries was sent for sequencing by SeqMatic (Fremont, CA). Sequencing was performed using a MiSeq V2 300-cycles format (Illumina, Inc.). All bioinformatic analyses were performed on KBase (Arkin et al. 2018). Raw reads (.fastq) were analyzed with FastQC, revealing the presence of Nextera Transposase Sequences on some reads. Reads were trimmed of adaptors and low-quality reads with Trim Reads with Trimmomatic - v0.36 (Bolger et al. 2014) set to remove NexteraPE-PE adaptors. Quast v4.4 (Gurevich et al. 2013) was used to assess whether SPAdes v3.15.3 (Bankevich et al. 2012) or IDBA-UD v1.1.3 (Peng et al. 2012) assembled reads in a more complete manner. SPAdes v3.15.3 was chosen as it produced assemblies are composed of fewer contigs of larger N50 scores. The Illumina draft genomes yielded 83-281 contigs (Table S1).

We sequenced SM743_UCD567 using an Oxford Nanopore sequencing service provided by Plasmidsaurus (Eugene, OR). The Qiagen DNeasy Blood and Tissue Kit (Qiagen, Germantown, MD) was used for genomic DNA extraction using the manufacturer’s protocol for gram-negative bacteria. Library prep, sequencing, and assembly were all carried out by Plasmidsaurus. To briefly describe their methods, library prep is performed using the v14 library prep chemistry developed by Oxford Nanopore Technologies (Oxford, UK). Sequencing was performed using the R10.4.1 pore on a PromethION flow cell. For assembly, the worst 5% of reads were removed using Filtlong v0.2.1 with default parameters (github.com/rrwick/Filtlong). An assembly was generated using Flye v2.9.1 (Kolmogorov et al. 2019) with parameters set for high quality Oxford Nanopore Technology reads. Genome annotation was performed using Bakta v1.6.1 (Schwengers et al. 2021); contig analysis was performed using Bandage v0.8.1 (Wick et al. 2015); genome completeness and contamination was checked using CheckM v1.2.2 (Parks et al. 2015); and species identification was performed using Mash v2.3 (Ondov et al. 2016), Sourmash v4.6.1 (Titus Brown and Irber 2016).

To check for contamination and completeness of assemblies, CheckM (v1.0.18 for all isolates except SM743_UCD567, v1.2.2 for SM743_UCD657) was used (Parks et al. 2015). The assemblies were more than 99% complete and had less than 1% contamination, so they were used for phylogenetic analysis.

### Phylogenetic tree from KBase

To build a phylogenetic tree of the 25 new genomes and 398 public genomes, we used the KBase app: Insert Set of Genomes Into SpeciesTree - v2.2.0 (Arkin et al. 2018). This KBase app creates a phylogenetic tree based on a multiple sequence alignment of 49 conserved COG gene families, and creates a tree using FastTree2 (Price et al. 2010). The .newick file was uploaded into iTol to annotate the tree and modify the aesthetics (Letunic and Bork 2021). The full-length tree is available on the FigShare Repository (doi.org/10.6084/m9.figshare.23733567).

### Endoglucanase (egl) gene sequence analysis

We extracted the partial *egl* sequences from the draft genomes to assign the isolates to sequevars (Fegan and Prior 2005). We used the KBase BlastNv2.13.0+ app to identify the *egl* genes in each genome. In order to export the gene sequences, we ran the MUSCLE v3.8.425 App, which allowed us to export the sequences in FASTA format. We used the recently published protocol (Cellier et al. 2023b) to correctly trim the sequences according to international references. Sequences were analyzed with Geneious Prime 2021.1.1 software (Kearse et al. 2012) and aligned, along with international references, through the MUSCLE algorithm (Edgar 2004). Phylogenetic tree reconstruction was performed using PhyML v3.3.20180621 (Guindon et al. 2010). The determination of sequevars was assumed by partial *egl* sequence divergence values less than or equal to 1% (Fegan and Prior 2005) and to international reference sequences (Cellier et al. 2023b).

### Host resistance screening

Seeds of 37 accessions of tomato, eggplant and pepper were obtained from AVRDC (The World Vegetable Center, Taiwan), INRAE (Institut National de Recherche pour l’Agriculture, l’Alimentation et l’Environnement, France), BARI (Bangladesh Agricultural Research Institute, Bangladesh), and Makerere University, Uganda (Table S2). The pedigree of 30 Core-TEP accessions is described by Lebeau et al. 2011. BARI 2 and BARI 8 are resistant tomato and eggplant lines, respectively, developed by BARI. Tomato MT56 was received from Uganda but its pedigree is uncertain. Eggplant EG190, EG219 and tomato BF Okitsu were developed by AVRDC. *S. sisymbriifolium* is a common weed in South Asia that is used as a bacterial wilt resistant rootstock in South Asia.

Seeds were sown in plastic trays with 2.5 x 2.5 cm^2^ cells containing planting medium (Sungro Horticulture, Agawam, MA). Four-week-old seedlings were soil-drench inoculated with a 5 ml suspension (1×10^8^ CFU/ml) of one of six RSSC isolates from eggplant, pepper, or grafted tomato or eggplant (SM701 (eggplant), SM716 (pepper), SM732 (eggplant grafted onto *S. sisymbriifolium*), SM738 (eggplant), SM743 (tomato grafted onto *S. sisymbriifolium*), and MB1 (eggplant)) selected based on host, origin, and genetic diversity, determined as described above. Inoculum was prepared in sterile distilled water from 48 hr-old cultures growing on casamino acid, peptone, glucose (CPG) agar at 28°C. Seedlings were inoculated following root wounding with a sterile scalpel blade. Wilt incidence was recorded twice weekly for 5 weeks after inoculation. The experiment was conducted twice as a randomized complete block design with three replications (blocked by time) of 15 plants per replication, with a split-plot arrangement. *R. pseudosolanacearum* isolates were applied as the main plot effect and seedlings were arranged in sub-plots.

## Results

### RSSC isolates in South Asia

We isolated 99 RSSC isolates from nine major solanaceous vegetable growing regions of Bangladesh and 20 RSSC isolates from five regions of Nepal. In Bangladesh, 77 isolates from eggplant, nine from grafted eggplant/*S. sisymbriifolium*, and 13 from pepper. In Nepal, ten isolates were recovered from eggplant, four from tomato, two from potato, and four from grafted tomato/*S. sisymbriifolium*. An additional six isolates were obtained from four states in India (Fig 1A and Table S1). The phylotype-specific multiplex PCR (Pmx-PCR) was applied on all isolates. All reactions yielded the RSSC-specific amplicon (282 bp) and the phylotype I-specific amplicon (144 bp), indicating that all isolates belong to the *R. pseudosolanacearum* species (Fegan and Prior 2005; Safni et al. 2014).

Although all newly described isolates in this study belong to phylotype I, it is known that other RSSC lineages are present in South Asia. We queried the Ralstonia Diversity Database version 4 (Lowe-Power et al. 2022) for isolates isolated in the South Asian countries: Bangladesh, Nepal, India, and Sri Lanka. RSSC isolates were frequently reported on potato (n=185), eggplant (n=95), tomato (n=42), pepper (n=33), and ginger (n=17) (Fig 1B). Of the 477 isolates reported in the literature, the phylotype was identified for 245 isolates (Cellier and Prior 2010; Ghorai et al. 2022; Ramesh et al. 2014; Gurjar et al. 2015; Sagar et al. 2014; Kumar et al. 2014; Patil et al. 2017; Cellier et al. 2012). In the available data, phylotype I accounts for all of the published reports of RSSC on eggplant, tomato, and pepper in South Asia. More RSSC lineages have been reported on potato in South Asia: the pandemic IIB-1 lineage (n=87), phylotype IV (n=29), and phylotype I (n=23).

### Genome analysis

We sequenced the genomes of 20 isolates from Bangladesh and five isolates from Nepal. We built a phylogenetic tree of these 25 isolates and 398 publicly available RSSC genomes (Fig 2 and Fig S1). Most of the Bangladesh and Nepal isolates (n=18 and n=3, respectively) clustered together in nine clonal groups on four major branches with phylotype I isolates from India (n=1), Sri Lanka (n=1), China (n=1), and Brazil (n=1). The Bangladesh isolates were isolated from eggplant (n=16) and pepper (n=2) in Bogra (n=2), Jamalpur (n=3), Brahmanbaria (n=2), Cumilla (n=4), Jashore (n=2), Joydebpur (n=1), and Narsingdi (n=4). The Nepal isolates were isolated from eggplant in Chitwan (n=3).

**Fig 2.**
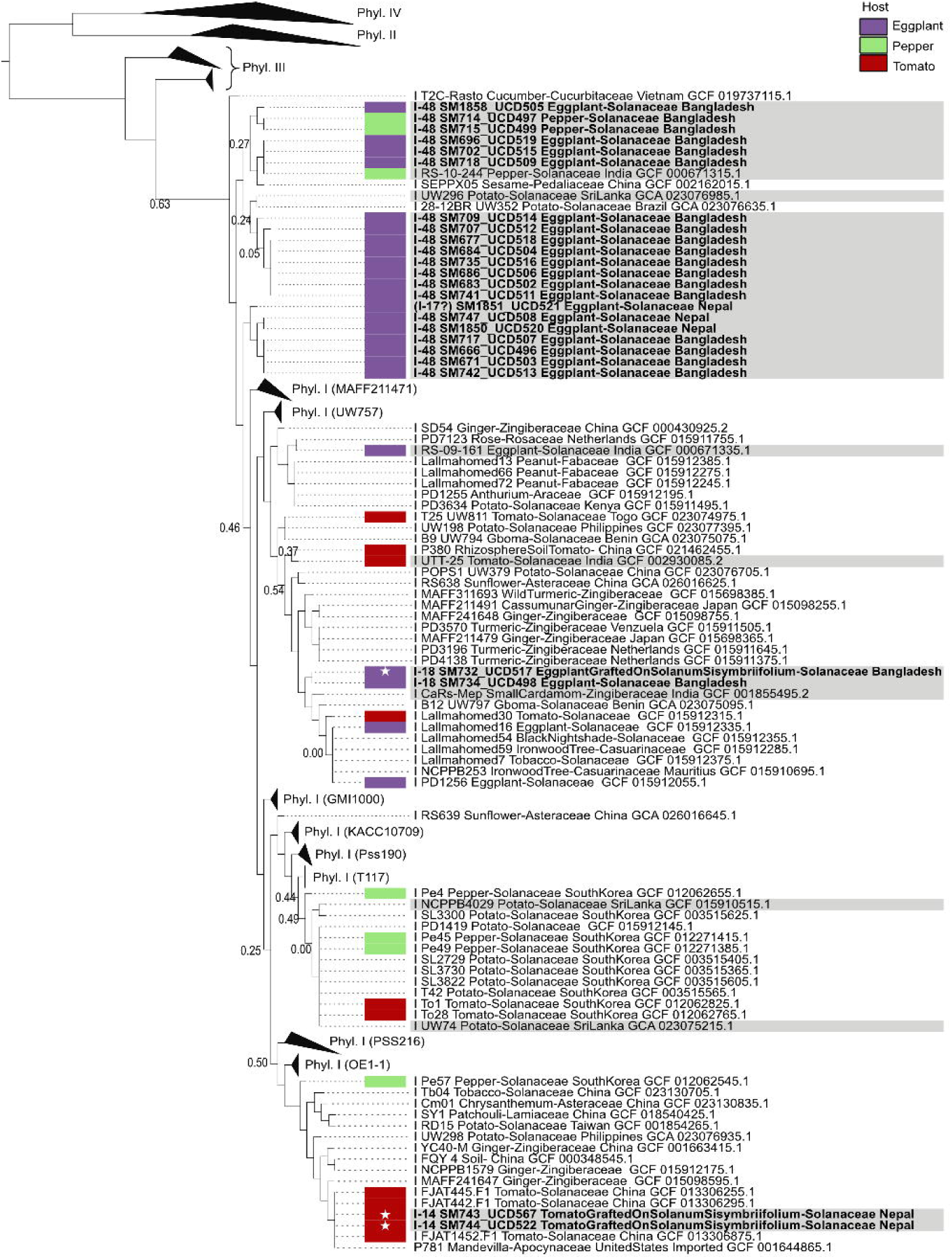
Phylogeny of South Asian and global RSSC. The phylogenetic tree was built with the KBase SpeciesTree tool, which creates a multiple sequence alignment of 49 conserved bacterial genes and generates a tree using FastTree. Analysis of the *egl* sequence suggested that the *R. pseudosolanacearum* genomes sequenced in this study belong to sequevar 48, 14, 18, and 17. Because the sole sequevar 17 assignment to SM1851 was incongruent with the KBase tree, we indicate uncertainty in this assignment with “(17?)”. Grey shading indicates isolates from South Asia. Bold indicates genomes sequenced in this study. Purple, red, and light green rectangles identify isolates isolated from eggplant, pepper, or tomato. White stars indicate isolates isolated from crop hosts that were grafted onto *Solanum sisymbriifolium* rootstock. Phylotype I clades without South Asian isolates were collapsed to triangles that reflect the amount of genetic diversity within the collapsed clade. Per triangle, one representative genome from the clade is listed. Additionally, phylotype II, III, and IV clades were collapsed. Bootstrap values are only listed if less than 0.70. A searchable PDF of the full tree is available on the FigShare repository (doi.org/10.6084/m9.figshare.23733567).

The remaining four isolates clustered in two distant branches. Two isolates isolated in Syangja, Nepal from tomato grafted onto *S. sisymbriifolium* (SM743 and SM744) formed a clonal group with three tomato isolates from China and an isolate from Mandevilla ornamentals imported into the U.S. The two isolates isolated in Tangail, Bangladesh from eggplant and eggplant grafted onto *S. sisymbriifolium* (SM734 and SM732, respectively) formed a clonal group that clustered close to isolates isolated from diverse locations (India, Benin, Mauritius, Japan, and unknown locations).

### Endoglucanase gene sequence analysis

We extracted the partial *egl* sequence from the genomes of the 25 isolates to assign these isolates to sequevars. The sequevars are listed on Fig 2 and the full *egl* tree is available on the FigShare Repository (doi.org/10.6084/m9.figshare.23733567). The *egl* tree and sequevar assignments were largely congruent. The majority of the isolates were assigned to sequevar 48, and the clonal SM743/744 isolates were assigned to sequevar 14. The clonal SM732/734 isolates were assigned to sequevar 18 although they have a relatively low whole-genome average nucleotide identity with the reference sequevar 18 isolate GMI1000 (estimated 98.80-98.88% by FastANI (Jain et al. 2018)). Based on *egl* sequence, SM1851 would be assigned to sequevar 17 even though it clusters within the 21 sequevar 48 isolates on the 49-gene tree (Fig 2).

### Host resistance phenotyping

We tested the resistance of 37 tomato, eggplant, pepper, and *S. sisymbriifolium* accessions against six South Asian isolates from distinct regions (Fig 3A and Table S2). The mean incidence of wilt in the susceptible tomato (L390), eggplant (MM136) and pepper (Yolo Wonder) controls was 85.5, 83.6 and 50%, respectively. Six tomato accessions demonstrated consistently high resistance (mean wilted plants ≤ 10% with no isolate causing > 20% wilting) against all six isolates: L285, Mt56, Hawaii 7996, CLN1463, TML46, and R3034. Four additional tomato accessions displayed a bimodal phenotype of susceptibility to SM738, MB1, and SM732 and resistance to SM743, SM716, and SM701: IRATL3, NC72 TR4-4, CRA66, and BF Okitsu. Bari2 was susceptible to SM738, MB1, and SM732, and moderately resistant to SM743, SM716, and SM701. In addition to L390, Okitsu Sozai no. 1 was highly susceptible to all isolates. Eight eggplant accessions displayed high resistance to all six isolates: Eg190, S56B, MM853, Bari8, MM643, EG203, MM152, and Eg219. Three accessions had moderate resistance: MM931, MM195, and MM960. In addition to MM136, MM738 was highly susceptible. Except for PM702, all pepper accessions were resistant to two of the isolates: MB1 and SM732. Three pepper accessions displayed high resistance: PBC631A, PBC66, and 0209-4. The responses of PM659 and PBC384 trended towards resistance. PM1022, PM1443, PM687, and Yolo Wonder were susceptible to the four pepper-virulent isolates. The *S. sisymbriifolium* accession displayed no symptoms after inoculation with four of the isolates, including SM732, which had been isolated from eggplant grafted to this rootstock. Isolates SM743, isolated from grafted tomato, and SM716, isolated from pepper, caused wilt incidences of 19.5% and 33.2%, respectively.

**Fig 3.**
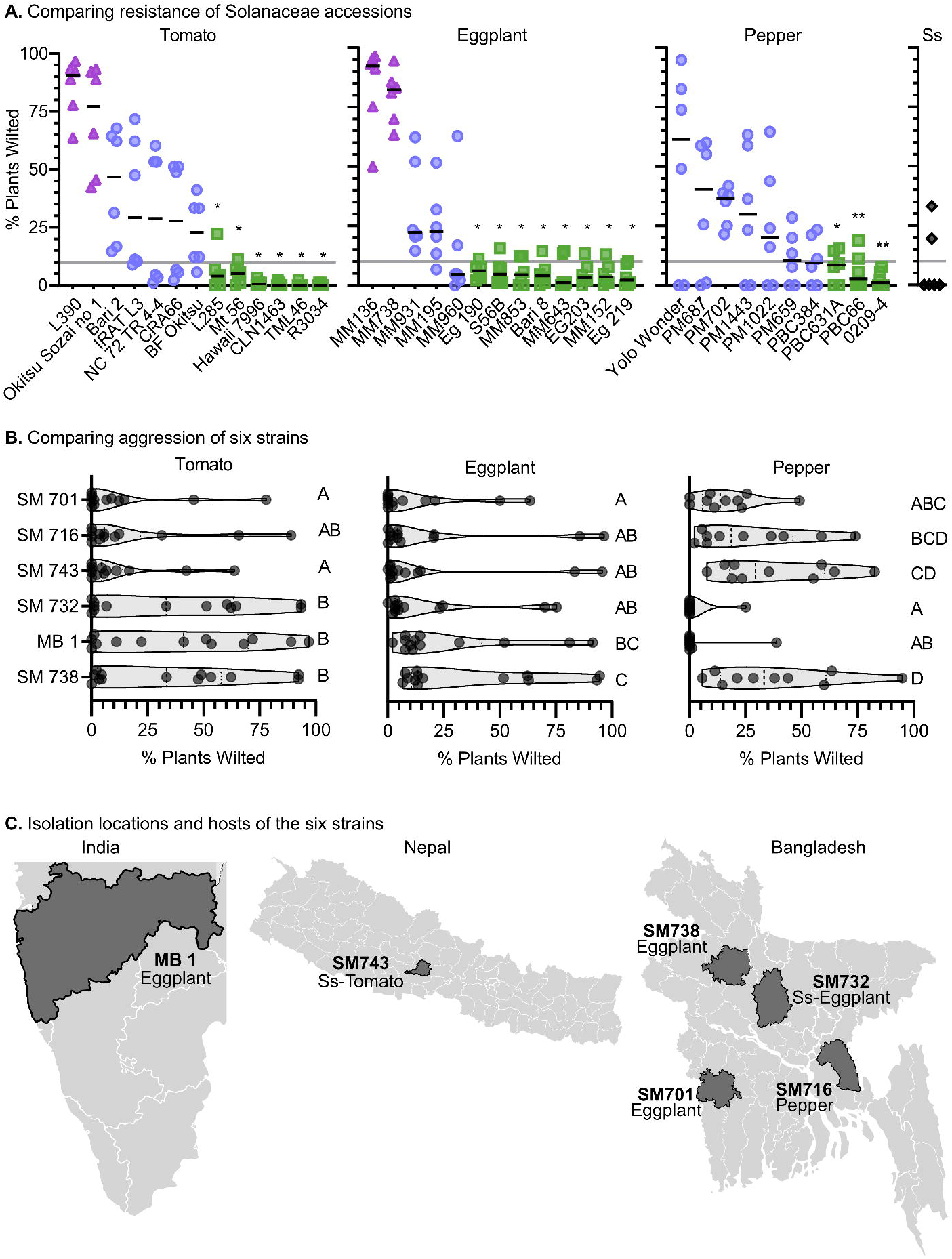
Disease interactions of 13 tomato accessions, 13 eggplant accessions, ten pepper accessions, and one *Solanum sisymbriifolium* (Ss) accession against six South Asian phylotype I isolates. Four-week-old seedlings were soil drench-inoculated with 5 ml of bacterial suspension (10^8^ CFU/ml) following root injury. The experiment was conducted twice as a randomized complete block design with three replications (blocked by time) of 15 plants per replication. Each point represents the average wilt incidence of two experiments recorded five weeks after inoculation. Isolates were SM701 (eggplant in Jessore, Bangladesh), SM716 (pepper in Comilla, Bangladesh), SM732 (eggplant grafted on *S. sisymbriifolium* in Tangail, Bangladesh), SM738 (eggplant in Bogra, Bangladesh), SM743 (tomato grafted on *S. sisymbriifolium* in Syangja, Nepal), and MB1 (eggplant in India). (*A*) Relative resistance of tomato, eggplant, and pepper accessions. Asterisks indicate significance compared to the most susceptible cultivar (L390 tomato, MM136 eggplant, and Yolo Wonder pepper) based on p<0.05 with Friedman test and Dunn’s multiple comparison correction. (*B*) Relative virulence of the six isolates across the accessions. Each symbol indicates the mean incidence of the isolate on a single accession. Letters indicate significance groups (p<0.05) by Friedman test and Dunn’s multiple comparison correction. (*C*) Origins of the six South Asian isolates.

### Comparative virulence of South Asian isolates

Aggressiveness of the *R. pseudosolanacearum* isolates varied with host species, and among accessions within a species (Fig 3B-C and Table S2). The SM738 isolate was the most aggressive, causing more than 20% wilt incidence on seven of 13 tomato, five of 13 eggplant, and seven of 10 pepper accessions. Isolates SM716, SM743, and SM701 displayed consistent patterns of virulence and wilted most pepper accessions. They had no-to-low virulence on tomato accessions, including five tomato accessions that were moderately susceptible to the other three isolates. Additionally, SM716 and SM743 were the only isolates that caused wilting in *S. sisymbriifolium*. Two isolates were largely non-pathogenic on pepper: MB1 and SM732. The genomes of SM732 and SM743 are sequenced. Unfortunately, as of 2022, stocks of the other four isolates were not culturable anymore under standard culture conditions so we were unable to sequence their genomes.

## Discussion

Bacterial wilt is one of the most important diseases of tomato, eggplant, and pepper in South Asia. This disease is difficult to manage due to the diversity, adaptability, and environmental survivability of the *Ralstonia* wilt pathogens. Host resistance is one of the best options available to manage this disease. However, the strain specificity of host resistance limits utility of this approach (Wang et al. 2013; Lebeau et al. 2011; Méline et al. 2023). Only pathogen-targeted management approaches, which require prior knowledge of local pathogen populations, can provide satisfactory and sustainable control of this disease. Therefore, we characterized the diversity of RSSC isolates collected from South Asia and screened a worldwide collection of resistant tomato, pepper, and eggplant accessions against representative South Asian isolates to identify suitable hosts that can potentially be used to manage bacterial wilt in the region.

Although several phylotypes are present in the region, all isolates in this study were identified as *R. pseudosolanacearum* phylotype I. It is possible that this outcome is because the majority of the isolates from this study were purified from wilted pepper and eggplant. Prior studies, including our meta-analysis of 8,000 RSSC isolations, have shown that phylotype I isolates are the most common etiological agents of bacterial wilt on eggplant and pepper while all RSSC phylotypes are commonly isolated from tomato plants (Gurjar et al. 2015; Sagar et al. 2014; Ramesh et al. 2014; Kumar et al. 2014; Hossain et al. 2022; Lowe-Power et al. 2022). Globally, phylotypes II and III have both been occasionally isolated from eggplant and pepper (Cellier and Prior 2010; Ravelomanantsoa et al. 2016; Lee et al. 2020; Deberdt et al. 2014; Bihon et al. 2020; N’guessan et al. 2013; Sedighian et al. 2020; Safni et al. 2014), while phylotype IV has been isolated from pepper but has not been reported on eggplant (Safni et al. 2014). Including this study, phylotype I accounts for 92.8% and 90.4% of the global RSSC isolations on eggplant (n=446) and *Capsicum* sp. pepper (n=365), respectively. If we had collected more tomato and potato isolates, we may have found more phylotype II and IV isolates in our survey because these phylotypes are known to be present in the region on these crops. A survey for RSSC in potato growing regions of Bangladesh purified RSSC isolates of undetermined phylotype(s) in Jamalpur, Nilphamari, and Munshigonj, while the disease was not detected in four other states during that survey (Ahmed et al. 2013). Further work is needed to investigate *Ralstonia* diversity in the region.

Regardless of the original host, all six isolates tested in this study were highly virulent on wilt-susceptible tomato and eggplant accessions, while two of six isolates (from eggplant or grafted eggplant) were avirulent on the wilt-susceptible pepper variety Yolo Wonder. The remaining four isolates were highly virulent on this variety. The isolate SM743, originally isolated from a wilted tomato scion grafted onto *S. sisymbriifolium* rootstock, was highly or moderately virulent on two eggplant and five pepper accessions. This suggests that, despite one-third of the isolates tested being avirulent on all but one pepper accession, recommendations for crop rotations away from solanaceous species should be followed, particularly when wilt-susceptible varieties are deployed.

Based on the genome sequences from this study, genomes from other studies (Patil et al. 2017, 2020) and prior studies with single gene markers (Ramesh et al. 2014), it is clear that there is considerable diversity of phylotype I RSSC in South Asia, consistent with the theory that phylotype I originated in Asia (Villa et al. 2005). In addition to the diverse, presumably endemic population of phylotype I isolates, we identified at least two lineages that may have been more recently introduced to Nepal and Bangladesh: SM743/744 and SM732/734, respectively. Isolates from these two genetically distant lineages were isolated from crops grafted onto *S. sisymbriifolium* rootstocks, and we confirmed that one isolate (SM743) caused wilting of *S. sisymbriifolium* in our greenhouse trial. There are anecdotal reports that the *S. sisymbriifolium* rootstocks are no longer providing effective mitigation of bacterial wilt in some locations in Bangladesh and Nepal (Subedi 2015). It is plausible that the reason for the breakdown of this host resistance is that exotic lineages have been introduced, and those exotic lineages happen to have genotypes that evade the immune surveillance of *S. sisymbriifolium*. However, the sample size of our study is too small to robustly test this hypothesis. Further studies are needed to understand the epidemiology of bacterial wilt in the region.

Sanger sequencing of a portion of the *egl* marker gene remains a popular way to classify isolates into sequevars based on the sequences. *egl*-based diversity analyses of phylotype I isolates should be treated with caution because there are instances where *egl* trees are incongruent with analyses using multiple genetic markers (Cellier et al. 2023a; Sharma et al. 2022; Rasoamanana et al. 2020). Because the *egl* trees rely on a short sequence, impeccable sequence quality and consistent methodology are essential for generating trustworthy conclusions. Here we compared our isolates to the established reference sequences for sequevars and used the recommended analytical methods (Cellier et al. 2023b). This allowed us to confidently assign sequevar 48 to 20 genomes, sequevar 14 to two genomes (SM743/744), and sequevar 18 to two genomes (SM732/734). We identified one conflict case (SM1851) where sequevar assignments based on the *egl* marker contradicted the position of the genome in the 49-gene tree. Hence, we have low confidence when assigning SM1851 into sequevar 17, knowing that our prior analysis also indicated that this sequevar has a polyphyletic nature within phylotype I (Sharma et al. 2022).

Due to the decade-long time frame of this study, we used classical and contemporary methods to characterize diversity of RSSC isolates from South Asia. At the time this study was initiated, the biovar system and genomic fingerprinting were common methods for RSSC diversity studies (Fonseca et al. 2014; Lewis Ivey et al. 2007; Norman et al. 2009; Xue et al. 2011; Zulperi et al. 2014; Ramsubhag et al. 2012). However, fingerprinting profiles cannot be compared between laboratories, which inhibits the utility of this approach to compare RSSC populations with published data. Consistent with the current paradigm, we found that both biovar and Rep-PCR classifications (data not shown) were discordant with phylogenetic clustering based on DNA sequence data. Similar inadequacies of Rep-PCR fingerprinting were recently reported for analyzing diversity of a different set of RSSC isolates from Bangladesh (Hossain et al. 2022). Currently, neither the biovar nor DNA fingerprinting is recommended for RSSC diversity analyses.

For RSSC diversity studies, we recommend always assigning the phylotype with the multiplex Pmx-PCR to all isolates. For more detailed analysis of RSSC diversity, we recommend *egl* sequence analysis according to the standardized protocol (Cellier et al. 2023b), using schemes with validated discriminating power (e.g. the RS1-MLVA13 scheme from (Cellier et al. 2023a)), or using whole genome analysis. Of these technologies, RS1-MLVA13 is best suited for phylotype I epidemiological studies because it has a demonstrably high discriminatory power that is sufficiently cost-effective to be applied to the large numbers of isolates and enable meaningful and thorough epidemiological surveys (Cellier et al. 2023a).

Host resistance to bacterial wilt is quantitative, polygenic, strain-specific, and greatly influenced by the environment, including temperature, soil moisture, and pH (Acosta 1978; Hanson et al. 1996; Scott et al. 2005; Wang et al. 2013). Resistance against all bacterial wilt pathogens is unlikely to be bred or engineered into solanaceous hosts due to the high genetic diversity of RSSC pathogens. For example, most of the tomato accession Hawaii 7996’s quantitative trait loci for bacterial wilt resistance are strain-specific (Wang et al. 2013; Carmeille et al. 2006; Danesh et al. 1994; Mangin et al. 1999; Shin et al. 2020; Méline et al. 2023). Variation in RSSC host range is very common because each isolate wields 60-80 plant-manipulating effectors, and fewer than 10 effectors are broadly conserved among diverse RSSC isolates (Landry et al. 2020). Nevertheless, host resistance can be a part of effective, integrated bacterial wilt management because RSSC isolates are slow to spread to new locations in the absence of human-mediated movement of infected plant material. Thus, once it is possible to predict pathogen host range based on genomic sequence, it could be possible to deploy targeted host resistance based on knowledge of the RSSC genotypes in different regions.

An objective of AVRDC’s research on bacterial wilt resistance was to develop resistant lines with more than 90% survival rate (Hanson et al. 1996). With this framework, we identified 18 accessions with less than 10% wilt incidence to at least one RSSC isolate. However, among these 18 accessions, only one-third were highly resistant (≤ 10% wilting) to all six isolates: three tomato accessions (CLN1463, TML46, and R3034) in addition to the reference resistant line Hawaii 7996, one eggplant accession (EG219), and one pepper accession (0209-04). Among the INRAE accessions, three tomato accessions (CLN1463, TML46, and R3034) and no eggplant or pepper accessions were highly resistant all six isolates. Neither BARI accession nor the Mt56 accession were highly resistant to all isolates. In addition to breeding lines with polygenic bacterial wilt resistance, there is considerable promise in using transgenic approaches to move immune receptors from diverse plant species into crops. For example, transgenic tomatoes and potatoes expressing Efr, a pattern recognition receptor from *Arabidopsis thaliana*, demonstrate bacterial wilt resistance in field and greenhouse trials (Lacombe et al. 2010; Boschi et al. 2017; Kunwar et al. 2018). Additionally, cytoplasmic immune receptors like ZAR1 and Ptr1 can recognize effectors from some RSSC isolates and other pathogens (Ahn et al. 2023), leading to interest in transforming tomato and eggplant with Ptr1 to better manage bacterial wilt with host resistance (Haefner et al. 2023).

To effectively manage bacterial wilt with host resistance, there is a need for large-scale research that identifies the geographic distributions of RSSC genotypes and statistical/artificial intelligence models to predict host range from RSSC genotype. To reach these goals, funding is needed for (1) epidemiological surveys of pathogen populations in different regions, (2) quantitatively comparisons of disease outcomes with diverse pairings of host genotypes vs. pathogen genotypes, and (3) generation of host and pathogen genomic data to allow functional genomics and population genomics studies. Genomic data and phenotypic data should be published in both summarized and raw formats to make the data most valuable for future meta-analysis. For this reason, we recently published raw host-range and whole genome sequence data on 19 phylotype IIB-4 RSSC isolates (Beutler et al. 2022). In this study, we quantified wilt incidence on a panel of Solanaceae accessions that have previously been phenotyped against 12 global RSSC isolates and six RSSC isolates from Louisiana, U.S (Lebeau et al. 2011; Lewis Ivey et al. 2021). Across these studies, genomes are available for eight out of 24 *Ralstonia* isolates. Unfortunately, genomes cannot be sequenced for 14 of the phenotyped isolates, including four isolates from this study, due to a combination of regulatory hurdles and lost viability of stocks.

Overall, we characterized the diversity of RSSC isolates from solanaceous hosts in South Asia and identified the most resistant tomato, eggplant and pepper accessions that can potentially be used to manage bacterial wilt in South Asia. As the resistance of these tomato, eggplant, and pepper accessions were evaluated under greenhouse conditions in Ohio, U.S., they must be assessed in field conditions of South Asia before employing them at large scale. This study contributes valuable knowledge on the genetic diversity and host range of RSSC populations infecting solanaceous hosts in Bangladesh and Nepal.

## Data availability statement

The genome data are available on NCBI Assembly and NCBI SRA under BioProject PRJNA989236. Isolates that are indicated as viable in Table S1 are available from the Lowe-Power lab. High-resolution PDF format figures are available on FigShare (doi.org/10.6084/m9.figshare.23733567).

## Acknowledgements

We thank Dr. Jaw-Fen Wang, AVRDC, Taiwan, Dr. Marie Christine Daunay, INRAE, France, and Dr. Yousouf Mian, BARI, Bangladesh, for providing seeds for this study; and BARI and Nepal Agricultural Research Council (NARC) for providing laboratory facilities in Bangladesh and Nepal, respectively. This work was supported by the Agriculture Office within the Bureau for Economic Growth, Agriculture, and Trade (EGAT) of the U.S. Agency for International Development, under the terms of the IPM-CRSP (Award EPP-A-00-04-00016-00). We thank Gauri Achari, ICAR Research Complex for Goa, India for providing the Indian isolates and Caitilyn Allen, University of Wisconsin, for providing genomic DNA of reference isolates GMI1000, K60, UW386, and UW443. We thank Kimberly Grulla, Kasey Miqueo, and Su Tun (UC Davis) for technical assistance with DNA extractions and preliminary analysis of Illumina sequencing data. We thank Angela Nanes (OSU) for technical assistance.

## Figures and Tables

**Fig S1.**
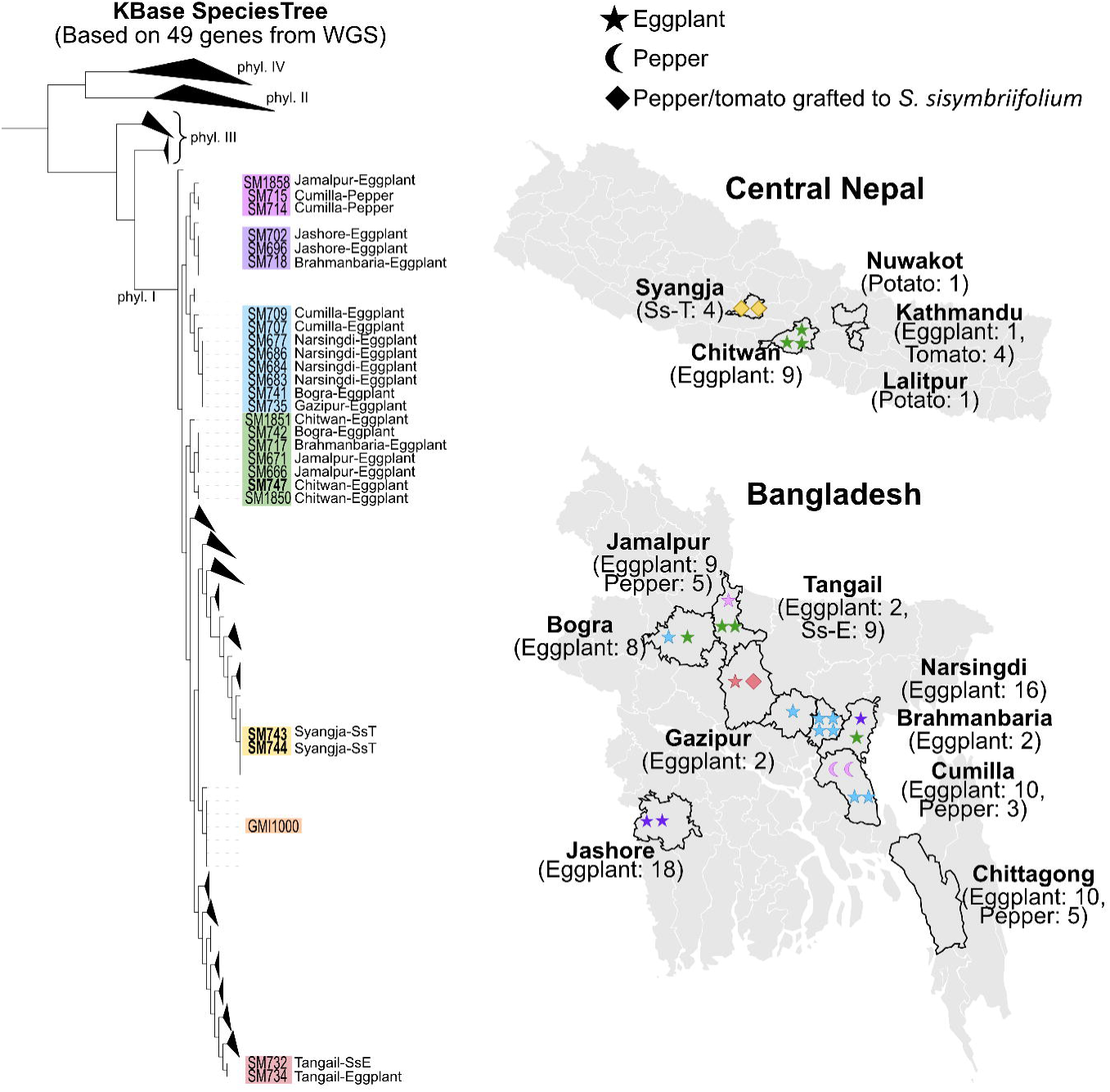
Comparison of phylogeny, isolation location, and host for the isolates with genomes sequenced in this study. Regions from which RSSC isolates originated from in this study are shown with black boundaries. The number of genomes sequenced from each region are shown with symbols, based on the host of isolation (eggplant, star; pepper, crescent moon; crops grafted to *Solanum sisymbriifolium*, diamond). Colors of symbols correspond to the clades as shown on the phylogenetic tree. Ss-T = tomato grafted onto *S. sisymbriifolium*; Ss-E = eggplant grafted onto *Solanum sisymbriifolium*.

